# Variation in the embryonic cell lineages of *Caenorhabditis elegans* natural populations

**DOI:** 10.64898/2026.11.03.764073

**Authors:** Charlotte Weymer, Ryan L Johnson, Jane Olivia Helou, Eric J Deeds, Erik C Andersen, Pavak K Shah

**Affiliations:** Bioinformatics Interdepartmental Program, University of California Los Angeles, Los Angeles CA 90095 USA; Molecular Biology Interdepartmental Program, University of California Los Angeles, Los Angeles CA 90095 USA; Molecular, Cell and Developmental Biology, University of California Los Angeles, Los Angeles CA 90095 USA; Department of Integrative Biology and Physiology, University of California, Los Angeles; Institute for Quantitative and Computational Biosciences, University of California Los Angeles, Los Angeles CA 90095 USA; Department of Biology, Johns Hopkins University, Baltimore MD 21218 USA

## Abstract

Developmental variation has often been appreciated as a major phenotypic substrate of evolutionary innovation in multicellular organisms. Although quantification of developmental variation in isogenic populations has a long history, it has been difficult to accomplish across natural populations until recently. Given its invariant cell lineage, *Caenorhabditis elegans* represents an ideal model system for studying how natural diversity in development is expressed at the resolution of single cells, cell lineages, and the whole embryo. Using embGAN, a deep-learning pipeline that enables label-free lineage tracing without the bottleneck and potential confounding effects of generating transgenic strains, we reconstructed the early embryonic lineage of 78 embryos spanning eight genetic backgrounds, including six wild strains, and quantified cell-cycle timing with a lineage-aligned graph metric that separates global rescaling of developmental rate from localized changes in lineage structure. We found that cell-cycle timing differs reproducibly among genotypes, with between-strain variation consistently exceeding within-strain variation. The magnitude of this variation was itself genotype-specific and not correlated with levels of genetic divergence. Unlike the global shifts in the tempo of development that dominate divergence between species, these differences are predominantly in lineage patterning, changing the timing of specific divisions, and they align to cell fate. Thus, the invariant lineage of *C. elegans* is built on a structured, genotype-specific landscape of developmental variation, a potential source of trait variation from which drift in developmental systems can emerge.

## Introduction

Multicellular development produces reproducible outcomes, yet quantitative variation in the timing and sequence of cellular events is pervasive, arising even among genetically identical individuals raised under controlled conditions^1–3^. Quantitative studies of development have characterized the magnitude of variation in development^4–7^, but how much variation is structured by natural variation in genotype vs environment and how that variation is distributed across the developing embryo remains sparsely characterized in any system, especially at the resolution of individual cells. Developmental systems drift, in which conserved developmental outcomes are produced by molecular programs that have themselves diverged extensively^8^, as has been demonstrated across taxa^9,10^, could play a role in shaping this type of variation.

Between related species, much of the difference in developmental rate has been shown to reflect a global rescaling of tempo, an approximately uniform acceleration or deceleration of an otherwise stereotyped program that arises from cell-intrinsic differences in biochemical reaction rates, protein stability, and metabolic flux rather than from a rearrangement of the developmental program itself^11–14^. It is not known whether genetic variation segregating within a species perturbs timing in this same global fashion or instead through localized, lineage-specific changes in the pacing of development. Distinguishing these possibilities requires separating uniform, scaling-based changes in the pace of development from relative changes in cell-cycle length specific to particular cells or sublineages, while accounting for trivial biophysical sources of variation that might confound these observations. Because many fates are specified not autonomously but through stereotyped inductive signaling^15,16^, structured variation is likely to be distributed across the lineage in a manner that results in lineage-specific patterns.

The genetic diversity now available in *C. elegans* strain collections enables us to interrogate the structure of developmental timing variation across genotypes^17^. Wild strains harbor punctuated, hyper-divergent regions that span roughly a fifth of the reference genome and carry haplotypes whose divergence rivals that separating distinct *Caenorhabditis* species, maintained by long-term balancing selection^18,19^. Across species, distinct pools of embryonic progenitors express consistent differences in cell-cycle timing^20–24^, suggesting that cell lineage-specific regulation of developmental tempo and cell-cycle timing might also vary across genotypes within a species. Although interindividual variability in *C. elegans* embryonic cell-cycle timing has been extensively characterized, aided by automated lineage tracing and the invariant lineage, it has been generally restricted to the analysis of variation within a single genetic background and its mutants^4,5,25,26^. We recently developed embGAN^27^, a deep-learning tool that allows label-free differential interference contrast (DIC) imaging to drive existing semi-automated lineage tracers^25^, removing the throughput ceiling of manual DIC lineaging and the requirement and potential confounding effects of fluorescence-based tracing.

Combining embGAN with manual curation, we reconstructed the lineages of 78 embryos across eight genetic backgrounds including the laboratory reference strain N2, an N2-derived transgenic line, and six genetically divergent wild strains. For each pair of embryos across this set, we quantified differences in cell-cycle timing with a lineage-aligned graph metric that separates a global rescaling of developmental rate from localized, structural changes in the shape of the lineage tree^28^. We found that cell-cycle timing differs reproducibly among genotypes, with variation between strains generally exceeding variation within them. The magnitude of that variation is itself genotype-specific and dissociable from genomic divergence, as some wild strains develop far less consistently than N2, and divergent strains differ sharply in their within-strain consistency. Genotype-specific differences in cell-cycle timing are predominantly due to relative differences in the cell-cycle durations of distinct cells rather than a uniform rescaling of the developmental clock, the opposite of the global tempo shifts that dominate between species. They also manifest in a lineage-specific manner, with the least inter-strain variation observed in the anterior AB sublineages. The invariant lineage thus overlies a structured, genotype-specific landscape of developmental timing variation, one that is compatible with the intraspecific origins of developmental systems drift and that offers a tractable entry point for dissecting the genetic basis of developmental precision.

## Results

### Selection of highly divergent wild strains

To sample the extent of natural variation in embryonic development, we selected six genetically divergent wild *C. elegans* strains from the *Caenorhabditis* Natural Diversity Resource (CaeNDR)^17^: CB4856, DL238, XZ1516, MY23, JU2001, and ECA36. These strains were chosen to span the major phylogeographic structure of the species, including strains from the Hawaiian Islands (CB4856, DL238, XZ1516) and New Zealand (ECA36), a region known to harbor disproportionate genetic diversity attributable to the species’ inferred Pacific origin^19,29^, alongside strains from Europe (MY23, JU2001). Recent population genetics showed that variation across the species is concentrated in punctuated hyper-divergent regions covering approximately 20% of the *C. elegans* reference genome, with haplotypes maintained by long-term balancing selection at levels of divergence comparable to that between *Caenorhabditis* species that diverged tens of millions of years ago^30^. Each strain in our panel carries a broad cross-section of the genomic diversity that has been mapped in these regions, ensuring that the panel samples not only common segregating variation but also the deep allelic divergence that distinguishes wild strains from the N2 laboratory reference background and from each other. Prior characterizations of natural variation in these strains have focused primarily on post-embryonic traits: growth rate, behavior, and susceptibility to drugs and toxicants^18^. We compared this panel against two reference backgrounds: the laboratory strain N2 and JIM113, an N2-derived transgenic line previously used to characterize embryogenesis under genetic and environmental perturbation^5,31^. To minimize extrinsic contributions to non-genetic variation, we sampled embryos exclusively from fed hermaphrodites and conducted all imaging in a temperature-regulated microscopy suite.

### Global and lineage-specific patterns of temporal variation

To examine whether differences in the duration of each cell’s cycle existed between strains, we computed the normalized deviation in cell-cycle duration for each cell in each embryo, defined as the difference between the mean for that cell type and the observed individual duration, divided by that mean; positive values thus correspond to faster-than-average cell cycles for that cell type (Figure 1A). Although most individual cells deviate from their mean in the dataset by less than 20%, several strains displayed consistent, lineage-wide shifts. For example, JU2001 exhibited a globally accelerated pace, with many of its cell cycles deviating 5-20% faster than the panel-wide average. Beyond such uniform shifts in global pace, individual lineages and cells deviated from the dataset mean in different ways. The endodermal E lineage was particularly notable in this regard, showing the largest magnitude of strain-specific effects in XZ1516 and JU2001.

**Figure 1:**
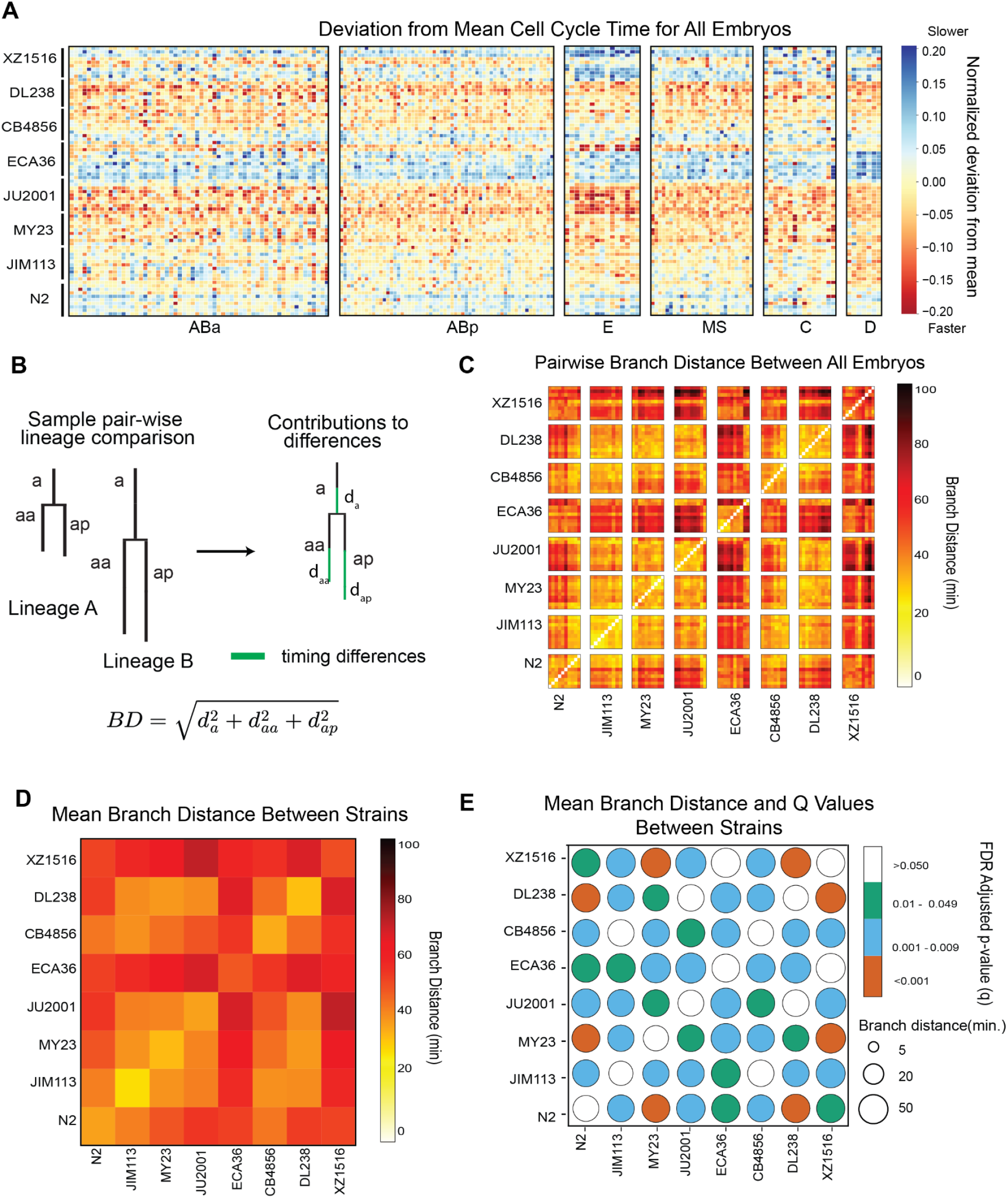
Natural variation in cell-cycle timing during *C. elegans* early embryogenesis. A: Heatmap showing normalized deviation from the cell-cycle dataset on a per cell basis for all embryos (n=77 embryos). Mean cell-cycle is calculated for the same cell across all embryos, and deviation from mean is calculated by subtracting a given cell-cycle from its mean and dividing by that cell cycle’s mean. B: Sample branch distance calculation given two lineages. The branch distance is calculated as the Euclidean norm on the cell-cycle times between matching cell identities between two embryos. C: Heatmap showing pairwise branch distance calculations between all embryos in the dataset, grouped by strain (n=77 embryos) D: Heatmap showing branch distance calculations averaged across all pairwise embryo calculations in each strain (n=77 embryos) E: Permutation test results on inter-strain cell-cycle timing distributions, colored by significance threshold and sized by inter-strain branch distance value.

To summarize lineage-aligned variation in cell-cycle duration, we used the branch distance^28^, a graph metric that we previously developed to quantify phenotypic differences mapped across cell-lineage traits. For any given pair of lineages, the branch distance is defined as the Euclidean norm on the vector of cell-cycle times between matching cells in each lineage being compared (Figure 1B), serving as a measure of dissimilarity in cell-cycle timing between a pair of lineages. We computed pairwise branch distances between every embryo in our panel (Figure 1C) and subsequently averaged these values on a per strain basis (Figure 1D). Within-strain distances were typically smaller than between-strain distances, indicating genetic influence on the timing of cell divisions. Notably, N2 and JIM113 embryos showed dramatically distinct magnitudes of within-strain variation with a larger mean inter-strain distance than intra-strain, indicating that the integrated transgenic array in JIM113 measurably perturbs lineage timing relative to its N2 parent, as we previously observed^28^.

The XZ1516 strain breaks the general trend with within-strain noise exceeding the magnitude of many between-strain differences. This observation has two important implications. First, it suggests that XZ1516 embryos have a reduced level of cell-cycle timing precision relative to other genetic backgrounds, hinting at strain-specific differences in developmental precision. Second, it cautions against interpreting strain-level mean phenotypes in isolation because the within-strain variance structure of a trait may be as informative as the population mean for understanding how a specific genotype shapes embryogenesis. To formally test whether apparent grouping of embryos by strain reflected strain-specific lineage structure rather than chance, we performed a permutation analysis in which strain labels were randomly shuffled across embryos and pairwise inter-strain differences were compared to within-strain differences under the permuted null. We found that nearly all inter-strain comparisons were significant within a false discovery rate-corrected p-value threshold of 0.05, with the sole exceptions being CB4856 vs. JIM113, DL238 vs. JU2001, and XZ1516 vs. ECA36 (Figure 1E). The observed whole-embryo differences predominantly emerge in a genotype-specific manner. We sought to further understand these genotype-specific timing profiles by examining the within-strain variation and potential causes of this variation.

### Quantifying intra-strain developmental variation

Measures of intra-strain variation were calculated by computing the branch distance amongst all embryos within a given strain. From this analysis, we observed a substantial range of within-strain consistency across the panel (Figure 2A). JIM113 exhibited the lowest intra-strain variation with a median intra-strain branch distance of 34.1 minutes and a narrow IQR of 6.8 minutes. DL238 displayed a comparably tight distribution with a median intra-strain branch distance of 38.6 minutes and an IQR of 5.0 minutes. Two strains stand out with elevated within-strain variation: ECA36, with a median intra-strain branch distance of 47.5 minutes and the largest IQR of 35.4 minutes and a conspicuously bimodal distribution of pairwise distances; and XZ1516 with a median intra-strain distance of 56.5 minutes and an IQR of 13.5 minutes. The bimodality of ECA36 can be attributed to two embryos, both imaged in the same session, that differ from the other embryos primarily based on an overall faster global rate that was particularly fast in the E lineage. As such, we cannot rule out an extrinsic source of variation confounding this observation.

**Figure 2:**
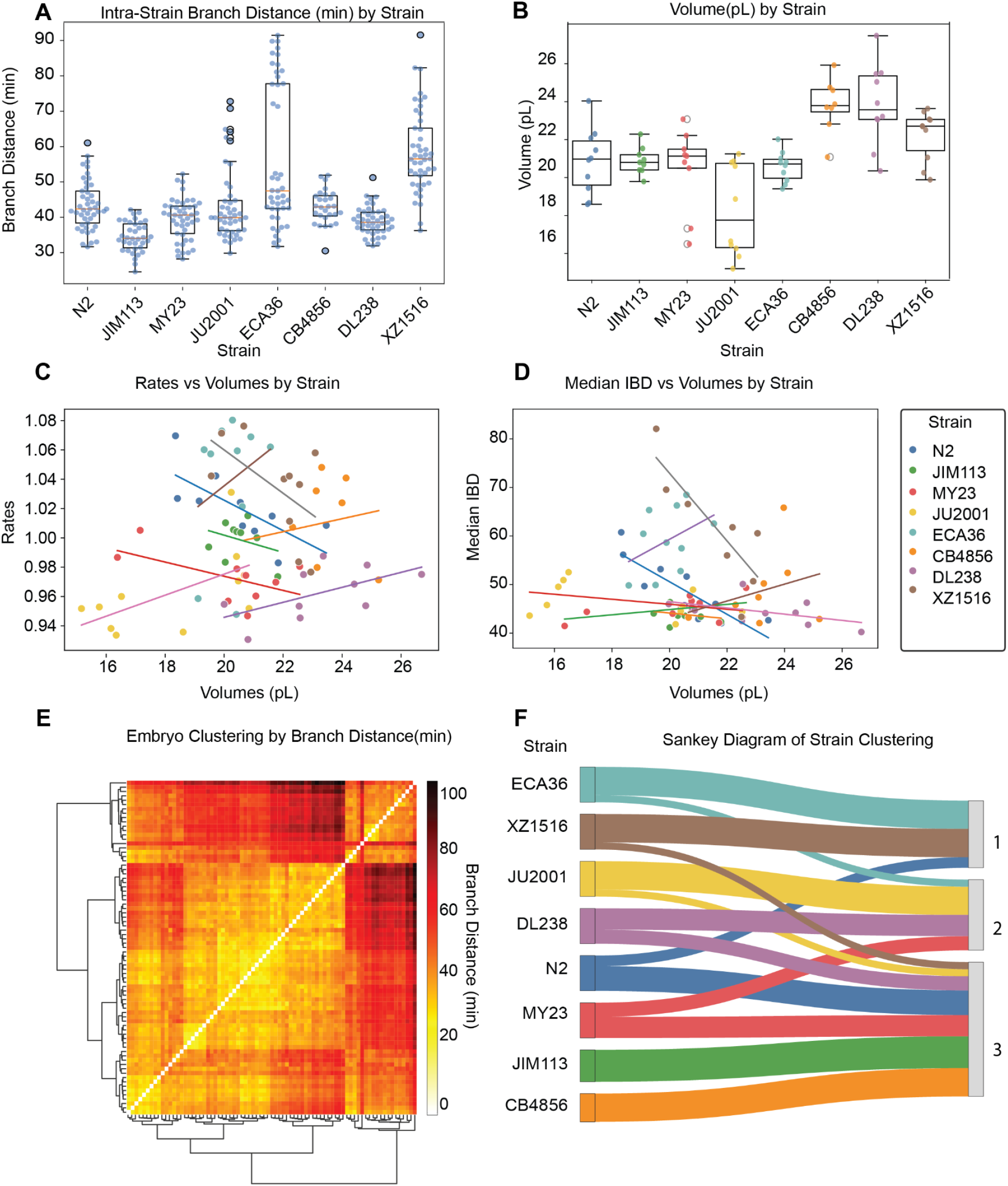
Intra-strain developmental variation. A: Intra-strain branch distance calculated in a pairwise manner between all embryos of the same strain. Each point represents an individual pairwise branch distance value, and box plot statistics were calculated on each strain. The red line represents the median value and the box bounds the 1st to 3rd quartiles. Whiskers extend to 1.5x the interquartile range (IQR), and black circles contain data points beyond these whiskers. B: Volume in pL calculated on all embryos and organized by strain. Volume was calculated using the SciPy convex hull function on the spatial coordinates of all nuclei for each embryo. C: Relative rate of development compared to the average embryo in the dataset plotted against calculated embryo volume. D: Median intersection branch distance plotted against calculated embryo volume. Median branch distance was calculated for each embryo by taking the median value for all its calculated pairwise branch distance values. E: Hierarchical clustering on branch distance matrix using Ward’s method on all embryos. F: Sankey diagram of strain mappings to the largest three clusters found from cutting the computed hierarchical dendrogram.

We next asked whether a simple biophysical mechanism (*i.e*., cell size) could account for these differences in within-strain consistency. Prior work has shown that cell and nuclear volumes in the *C. elegans* embryo obey a power-law relationship with cell-cycle duration, such that smaller cells cycle more slowly^32^. Under this model, intra-strain variability in embryo volume, and thus in the constituent cell volumes, could propagate into intra-strain variability in cell-cycle timing. We therefore tested whether strains with greater volume heterogeneity also exhibited greater timing heterogeneity and whether smaller or larger volumes correlated with global shifts in developmental tempo. We estimated embryo volume by computing the convex hull of all cell centroids detected over the duration of the lineage-tracing experiment. This measure systematically underestimates total embryo volume because the convex hull of nuclear centroids lies strictly within the eggshell and because mounted embryos are mildly compressed. We thus interpret volume comparisons in relative terms. Across the panel, estimated volumes ranged from 16 to 28 pL (Figure 2B). Despite this variation in volume, we found no consistent trend between the volume of an embryo and its overall rate of development, computed as the average relative change in cell-cycle length over all cells in the lineage relative to the dataset mean (Figure 2C) or median branch distance (Figure 2D). Indeed, the strain that shows the highest level of intra-strain heterogeneity in the average relative change in cell-cycle lengths, ECA36, exhibits the second lowest variation in embryo volume.

### Hierarchical clustering reveals strain-specific temporal identities

To resolve how cell-cycle timing differences reflect between and within-strain variation among the embryonic cell lineage, we performed hierarchical clustering on the branch distance matrix across all 78 embryos. The resulting dendrogram, which is shown along the axes of the reordered branch-distance heatmap (Figure 2E), revealed three prominent clusters. A Sankey diagram (Figure 2F) maps embryos from their strain of origin into the three clusters that were observed in the dendrogram. The reordered heatmap confirms that embryos within a cluster share more similar branch-distance profiles than embryos between clusters, as expected by definition. Importantly, embryos from the same strain did not always co-cluster, highlighting the intra-strain heterogeneity quantified above and indicating that strain identity is an imperfect predictor of any single embryo’s whole-lineage cell-cycle timing profile.

Because the great majority of embryo pairs in the dataset are between-strain pairs, the principal axes of variation captured by the clustering reflect overall conformity to, or divergence from, the dataset-wide mean rather than fine-grained strain-by-strain structure. Accordingly, the three clusters arrange themselves along a continuum of overall divergence: Cluster 1 contains the most divergent embryos, predominantly from ECA36, XZ1516, and N2; Clusters 2 and 3 contain progressively less divergent mixed groups drawn from many strains. Only two strains (JIM113 and CB4856) sort exclusively into a single cluster, indicating both high intra-strain consistency and close alignment with the bulk of the dataset. This result is surprising given that JIM113 is an N2-derived transgenic, yet diverges notably from its parental strain, and that CB4856 is a genetically divergent strain and carries many hyper-divergent haplotypes relative to the N2 reference^19,29^. That CB4856 nonetheless exhibits intra-strain consistency similar to N2, yet more homogeneously distributed among the population, shows that substantial genomic divergence does not by itself entail elevated developmental noise. The converse holds for XZ1516, another strain of Hawaiian origin that is more genetically hyper-divergent than CB4856 and is among the noisiest strains in our panel. Intra-strain robustness is thus dissociable from phylogeographic origin and instead points to specific genetic differences between strains.

The clustering does not fully recapitulate sorting by strain of origin of each embryo in the panel. Some N2 embryos cluster together with ECA36 despite limited genetic relatedness, and strains sharing qualitative degrees of hyper-divergence do not co-cluster as their shared population-genetic origin might naively predict. Although mean branch distance between strains generally exceeds within-strain distances, the distributions partially overlap (Figure S1) explaining how some embryos co-cluster with otherwise distinct strains. Branch distance-defined developmental identity is therefore neither a simple readout of genetic distance from the N2 reference nor of geographic origin. Together, these results indicate that strain background only partially constrains lineage-aligned timing: embryos from divergent strains can converge on similar lineage timing patterns, and embryos from a single strain can diverge to occupy distinct ones. These observations raise the question of where in the lineage strain-specific timing signatures arise, which we address by decomposing variation based on the lineage-specific founder-cell architecture of the early embryo.

### Strain-specific developmental rates are modulated by lineage-specific shifts in dynamics

To dissect the cellular origins of differences in cell-cycle timing between strains, we calculated normalized cell-cycle durations for each cell by dividing by the dataset-wide mean for each cell and averaged them within strains (Figure 3A). Here, deviations are reported as fold-change in cell-cycle duration, where values greater than 1 denote cells dividing more slowly than the panel-wide average and values less than 1 denote faster divisions. Most cells deviated by less than 15% from the dataset-wide mean, indicating high overall consistency in individual cell-cycle timing across strains. Nonetheless, several strains exhibited systematic shifts in developmental pace at the whole-embryo level. The two slowest strains, ECA36 and XZ1516, exhibited cell cycles 1.05 - 1.15× the dataset mean on an embryo-wide basis. Conversely, DL238 and JU2001 divided more rapidly, at 0.8 - 0.95× the cell-cycle duration of the average embryo.

**Figure 3:**
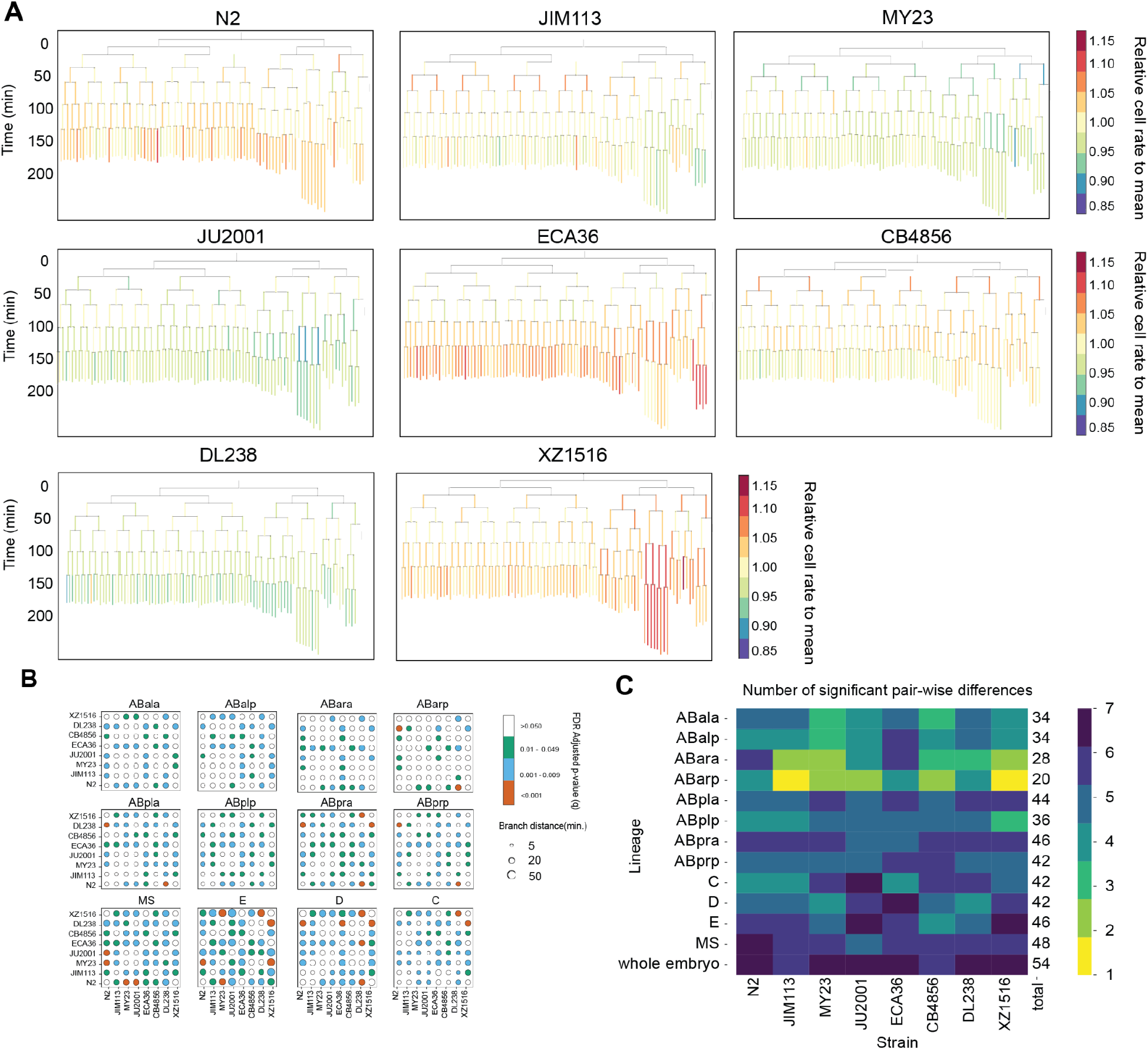
Inter-strain differences in developmental timing. A: Lineage diagrams showing relative cell-cycle rate to dataset-wide cell-cycle mean pooled amongst all embryos for a given strain. Relative rate is calculated on a per-cell basis by dividing a given cell-cycle time by the dataset cell-cycle mean. B: Permutation test results of inter-strain cell-cycle timing distributions decomposed by lineage, colored by significance threshold and sized by inter-strain branch distance value. C: Tabulated counts of significant pair-wise differences on the branch distance between strains at lineage resolution. Significance is thresholded by q<0.05 from permutation testing results.

Whole-embryo rate shifts, however, were not the only source of strain-specific deviation. Several lineages displayed timing shifts that diverged from, and in some cases exceeded, the average shift across the rest of the embryo. This shift was especially pronounced in the E lineage of two strains, the faster-than-average JU2001 and the slower-than-average XZ1516. In both cases, the E lineage exhibited timing shifts that were larger than the embryo-wide average, appearing especially fast in JU2001 and especially slow in XZ1516, as if exaggerating the underlying embryo-wide strain phenotype. Other lineage-specific deviations were also apparent. Most notably, the D lineage in ECA36 was markedly slow, with cell-cycle durations approximately 1.15× the dataset-wide mean and substantially longer than the embryo-wide ECA36 average. These observations motivated a systematic, lineage-resolved test of where in the embryo strain-specific timing differences were concentrated.

To formalize these observations, we asked which lineages contribute disproportionately to inter-strain differences. The whole-embryo branch distance summarized lineage-aligned variation but did not resolve which specific lineages could drive that variation. Therefore, we decomposed pairwise inter-strain comparisons by lineage and tested, for each lineage, whether observed timing differences between strains exceeded differences within strains using a permuted null in which strain labels were shuffled (Figure 3B). The L_2_-norm structure of the branch distance means that inter-strain variance is expected to grow additively as the number of cells within a lineage under comparison increases, so two factors govern a lineage’s baseline expected variability: the number of measured cells and the per-cell variance. These factors act in opposition in wild strain lineages. For example, the AB lineage contributes approximately ten-fold more cells, but the posterior lineages, with their longer cell cycles, contribute more absolute variance per cell under any noise model in which variance scales with mean duration. Under a constant-coefficient-of-variation null^31^, expected AB branch distances should exceed posterior lineage branch distances by approximately two-fold. Thus, a first-order expectation is that lineages with greater developmental depth and shorter constituent cell cycles should generate more variance, with the size of the disparity attenuated but not reversed by the longer cell cycles of the posterior lineages.

Our observations contradicted this expectation. Lineages of equivalent depth and similar mean cell-cycle durations differed markedly in inter-strain variability, and shallow lineages were not more consistent. ABa exhibited substantially higher similarity in inter-strain comparisons than ABp, as evidenced by its lower frequency of significant differences between strains by permutation testing (Figure 3B), despite having the same number of cells per lineage and similar cell-cycle durations. This asymmetry was striking given their shared ancestry. Their divergence into anterior (pharynx, anterior neurons) and posterior (hypodermis, posterior neurons, some mesoderm) fates is the primary difference between these lineages, downstream of Notch signaling^15^, suggesting that the difference in timing precision might reflect fate- or tissue-specific constraints on the temporal program rather than an inherent feature of larger lineages. Additionally, the posterior lineages (C and D) displayed an inter-strain difference frequency comparable to that of ABp. Considered together, the posterior lineages (ABp, MS, E, C and D) behaved in line with the expected behavior of the L_2_ norm, that inter-strain variability scales with the combined effects of cell count and per-cell-variance contributions from cell-cycle duration. ABa diverged from this observation, exhibiting markedly less inter-strain variation than either its sister lineage or the inherent scaling of the L_2_ norm would predict, suggesting a higher level of developmental precision in this lineage. Our permutation testing framework in comparing inter-strain distances to intra-strain distances inherently accounts for such differences in the magnitude of the branch distance independent of tree depth and mean cell-cycle length.

### Global rate and lineage pattern contribute to inter-embryo timing differences

Two distinct sources of variation became apparent from our analyses: differences in the global rate of development between two lineages and relative differences in the cycle lengths of individual cells within the lineages (Figure 4A). We controlled for differences in developmental rate caused by temperature^31^ by performing imaging in a climate-controlled space, meaning that observed rate differences are likely due to intra-strain variation or strain-specific rate differences. We therefore geometrically decomposed the branch distances into a parallel component (a measure of scaling) and a perpendicular component (a measure of relative differences in the lengths of distinct cells within the lineages) to determine the extent to which rate scaling and non-scaling changes contribute to the observed differences.

**Figure 4:**
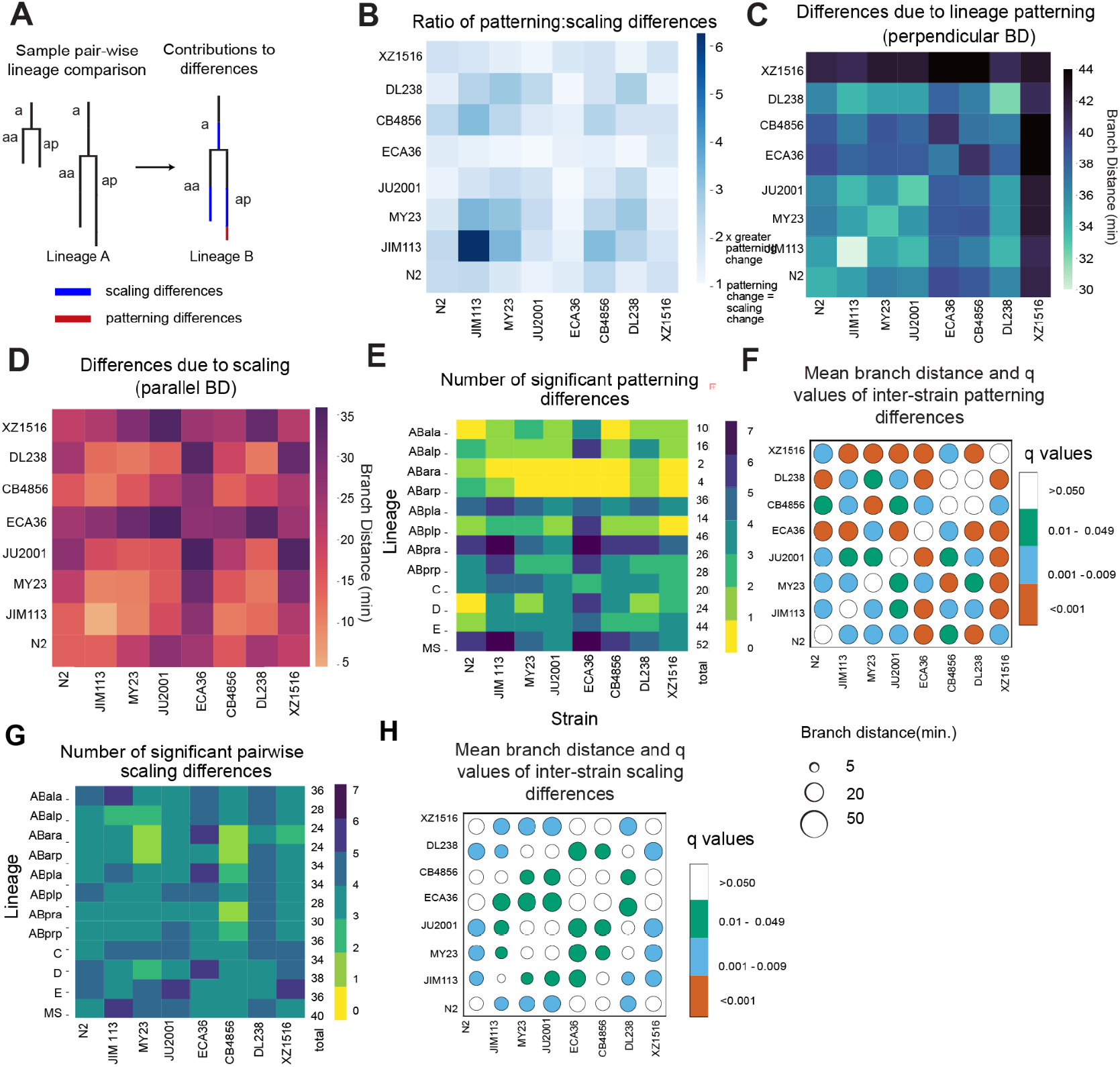
Global rate vs. patterning differences in lineage timing. A: Sample pairwise lineage comparison with both global rate and patterning differences. B: Visualization of the ratio of patterning differences to scaling differences on a per strain basis. This ratio is calculated by taking the pairwise perpendicular branch distance (patterning) divided by the parallel branch distance (rate). C: Heatmap of patterning differences on a pairwise strain by strain basis (decomposed perpendicular BD). D: Heatmap of scaling differences on a pairwise strain by strain basis (decomposed parallel BD). E: Tabulated counts of significant pairwise differences on the perpendicular branch distance between strains at lineage resolution. F: Permutation test results on whole-embryo inter-strain patterning differences(perpendicular BD), colored by significance threshold and sized by inter-strain branch distance value. G: Tabulated counts of significant pairwise differences on the parallel branch distance between strains at lineage resolution. H: Permutation test results on whole-embryo inter-strain scaling differences(parallel BD), colored by significance threshold and sized by inter-strain branch distance value.

We decomposed the intra-strain and inter-strain comparisons at the whole-embryo level into its parallel and perpendicular components, and found that the perpendicular component was larger than the parallel component in all cases (*i.e.*, the magnitude of patterning differences exceeded that of rate differences in all comparisons). Therefore, we visualized the ratio of the patterning difference to the rate differences in these comparisons (Figure 4B). The largest ratio was observed in the JIM113 intra-strain comparison where the relative differences were 6× the magnitude of the scaling differences. Although clock-rate differences exist between strains and even individual embryos within the same strain, their contribution to inter-strain differences is many times smaller than the relative differences in the cell cycles of individual cells in the lineages. Further visualization of these patterning differences shows that XZ1516 has particularly large relative lineage pattern differences compared to all other strains (Figure 4C). Additionally, both XZ1516 and ECA36 have comparatively large rate differences as compared to other strains (Figure 4D).

We conducted permutation testing on these results at both the lineage level and the whole-embryo level. Significant difference counts were found to be higher for scaling differences as compared to patterning differences at the lineage level (Figures 4E, 4G). However, at the whole-embryo scale, most strain pairs differed significantly in both components, with the perpendicular component producing more frequent significant differences (Figure 4F, 4H).

Where parallel component differences did emerge, they were enriched in specific strains. ECA36 exhibited strong scaling differences across multiple lineages and at the whole-embryo level, consistent with its globally slower developmental pace (Figure 4E, 4F). The rate-scaling differences of ECA36 were distributed across both its posterior and anterior sub-lineages, suggesting that the global slowdown observed in this strain reflects a coordinated rate shift on most of the embryo rather than a lineage-localized perturbation. The fewest inter-strain parallel component differences again emerged in ABa, consistent with the rate scaling conservation noted above.

Taken together, these results indicate that genetic differences between wild *C. elegans* strains manifest in early embryogenesis predominantly as non-scaling perturbations to the pattern of cell-cycle timing in the lineage rather than as uniform shifts in an embryo-wide developmental clock. Rate-scaling differences do exist, most prominently in ECA36 and XZ1516, but they are the exception across the embryo and the rarest source of significant variation, especially in ABa-derived lineages. ABa is thus highly precise along two axes: it is the most rate-consistent lineage across strains and the least variable in the pattern of cell-cycle timing among its cells. ABarp emerges as a sub-lineage of particularly high developmental precision. This observation is particularly striking since ABarp also exhibits the lowest level of left-right symmetry with any other portion of the AB lineage, giving rise primarily to hypodermal cells, sensory neurons in the head, and a left-right symmetric set of postembryonic blast cells (V1, V2, V4, V6). The remainder of the embryo, including ABp and the posterior lineages, accumulates strain-specific timing variation primarily through lineage-specific changes in the pattern of cell-cycle duration rather than coordinated shifts in a global rate of development.

## Discussion

The invariant cell lineage of *C. elegans* is a prototypical example of deterministic development, in which nearly every somatic cell can be assigned a unique identity and an invariant fate. We previously found that despite this cell-fate invariance, genotype-specific signatures of cell-cycle timing create structured patterns of variation in early embryogenesis^28^. By reconstructing the lineages of 78 embryos across eight genetic backgrounds and quantifying their differences at the single-cell, sublineage, and whole-lineage scale, we find that natural genetic variation pervades early embryogenesis as reproducible strain-dependent differences that are invisible at the level of cell fate but pronounced once timing is measured at single-cell resolution. Two features of strain-specific variation are apparent. First, the magnitude of variation is itself a genotype-specific property that differs between strains. Second, variation is predominantly in the pattern of cell-cycle times across the lineage, manifesting as localized changes in timing within particular lineages, rather than in a uniform rescaling of the developmental clock. Together, these findings establish cell-cycle timing variation in early lineage as a quantitative phenotype and a tractable entry point to dissect the genetic architecture of precision in metazoan development.

Decomposing the branch distance into a global scaling component and a perpendicular residual revealed that strain differences in cell-cycle dynamics are predominantly in the pattern of cell-cycle timing across the lineage rather than being attributable to global changes in tempo. This pattern stands in contrast with the interspecies literature, in which much of the divergence in developmental rate between related species reflects a global rescaling of tempo driven by cell-intrinsic differences in biochemical reaction speed, protein stability, and metabolism^12–14^.

One possible interpretation is that global tempo is relatively constrained within species, with most standing variation in a population manifesting as changes in cell-cycle timing at the resolution of individual cells or sub-lineages. By contrast, ECA36 exhibits differences in cell-cycle timing that are more evenly distributed as both mean rate shifts and single cell resolution variation. The localized change and rate-scaling axes thus appear genetically separable with lineage-autonomous regulators of cell-cycle progression potentially shaping the former and global determinants of developmental tempo likely shaping the latter. The separability of these modes of variation in our data suggests that they might be regulated by distinct classes of loci, a prediction that could be tested directly using quantitative trait mapping across an expanded panel of strains.

Variation is not uniformly distributed across the lineage, and its distribution is not fully explained by trivial properties of the metric or the lineage itself, such as the size of the lineage or the generation count of a cell. ABa and ABp arise symmetrically from the same parent, divide on similar timescales, and contribute an equal number of cells to the developmental window that we addressed, yet cell-cycle timing in ABa was markedly more consistent across strains than ABp in both rate scaling and the pattern of cell-cycle timing across the lineage. Because the principal systematic difference between these lineages is their downstream fates rather than any feature of their cell count or lineage topology, this asymmetry implicates fate and tissue-specific constraints on temporal variation, consistent with the view that the gene regulatory networks underlying particular fates could themselves be buffered to produce invariant and robust outputs^33^.

These patterns are unlikely to reflect simple biophysical scaling. Although cell-cycle duration in the early embryo scales with cell volume through a lineage-specific power law relationship^32^, we found no consistent relationship between estimated embryo volume and either developmental pace or intra-strain variability. ECA36, which exhibited the greatest heterogeneity in developmental timing, showed little variation in embryo volume, the opposite of what a cell size variance-driven model would predict. We interpret our volume estimates conservatively, because the convex hull of nuclear centroids systematically underestimates embryo volume and mounted embryos are mildly compressed, but the absence of any trend across strains argues that the timing variation that we observe arises from developmental processes rather than from systematic differences in cell size.

Our finding that within-species variation is predominantly in the pattern of cell-cycle times in the lineage rather than rate-scaling suggests that these two axes of developmental change could be modulated by distinct mechanisms and potentially on distinct evolutionary timescales, a pattern that comparative developmental timing data in vertebrate and invertebrate systems alike could test. The branch distance and its components are quantitative traits potentially well suited to genome-wide association and linkage mapping across the expanded panels of wild strains available through the *Caenorhabditis* Natural Diversity Resource^17^, an approach that could identify the loci tuning developmental tempo, structural timing, and developmental robustness. Further improvements to the automated performance of lineage tracing tools like embGAN will be necessary to support the throughput required to achieve this goal.

By quantifying development at single-cell resolution across a panel of wild strains, we have shown that the invariant lineage of *C. elegans* conceals a rich landscape of variation in the timing of divisions, organized primarily as lineage-specific structural rearrangements and varying in magnitude from one genotype to the next. The early lineage is a surface whose structured variation provides a tractable handle on the genetic architecture of developmental precision.

The analytical framework that we introduce here, combining label-free lineage tracing with graph-metric decomposition of timing variation, is in principle applicable to any organism whose early lineage can be resolved at single-cell level, offering a generalizable approach for mapping how genotype shapes the reproducibility of development.

## Methods

### Key resources table

| Reagent or Resource | Source | Identifier |
|---|---|---|
| **Bacterial and Virus Strains** |  |  |
| *Escherichia coli :* OP50 | Caenorhabditis Genetics Center | OP50 |
| **Experimental Models: Organisms/strains** |  |  |
| N2, *C. elegans* | Caenorhabditis Natural Diversity Resource | WB Strain: N2 |
| JIM113, *C. elegans* | Caenorhabditis Natural Diversity Resource | WB Strain: JIM113 |
| CB4856, *C. elegans* wild isolate | Caenorhabditis Natural Diversity Resource | WB Strain: CB4856 |
| JU2001, *C. elegans* wild isolate | Caenorhabditis Natural Diversity Resource | WB Strain: JU2001 |
| ECA36, *C. elegans* wild isolate | Caenorhabditis Natural Diversity Resource | WB Strain: ECA36 |
| DL238, *C. elegans* wild isolate | Caenorhabditis Natural Diversity Resource | WB Strain: DL238 |
| MY23, *C. elegans* wild isolate | Caenorhabditis Natural Diversity Resource | WB Strain: MY23 |
| XZ1516, *C. elegans* wild isolate | Caenorhabditis Natural Diversity Resource | WB Strain: XZ1516 |
| **Software and Algorithms** |  |  |
| Micro-Manager | GitHub | https://github.com/micro-manager |
| embGAN | GitHub | https://github.com/shahlab-ucla/embGAN |
| StarryNite | GitHub | https://github.com/zhirongbaolab/StarryNite |
| AceTree | GitHub | https://github.com/zhirongbaolab/AceTree |
| Adobe Illustrator | Adobe | https://www.adobe.com/products/illustrator.html |
| Python 3.11.12 | Python Software Foundation | https://www.python.org/downloads/release/python-31112/ |
| MATLAB R2022a | Mathworks | https://www.mathworks.com/help/install/ug/install-products-with-internet-connection.html |

### C. elegans strains

N2 reference and wild strains were sourced from the *Caenorhabditis* Natural Diversity Resource (CaeNDR). Strains were grown at 20℃ on nematode growth media (US Biological) and fed OP50 *E. coli* sourced from the *Caenorhabditis* Genetics Center (CGC).

### Imaging

Embryos were collected from gravid hermaphrodites picked from fed plates and cut in M9, releasing early stage embryos. Two- and four-cell embryos were collected by capillary aspiration and mounted between #1.5H coverslips with 20 μm diameter beads (Polysciences Inc.) and sealed with melted vaseline. No more than 2 embryos at these stages were observed per gravid hermaphrodite, although broodmates were not explicitly tracked during collection. Imaging was performed using an Olympus IX83 inverted frame equipped with a UPLSAPO60xs2 objective, ASI MX2000XYZ stage, and Hamamatsu Orca Fusion camera. Embryos were imaged every 60 s with a 750 nm z-spacing. DIC images were acquired with a 50 ms camera exposure, and the LED light source was tuned to not generate any saturated pixels in the image. DIC illumination was generated using an Olympus UCD8 manual condenser equipped with a U525 oil immersion 1.4 NA top lens and a DICTHR tilt-shift slider. The microscope room housing this instrument is maintained at 21℃ by forced air cooling, and temperature ranges in the room are checked daily using a thermometer that tracks min/max temperature ranges. Images were acquired using micro-manager and cropped and converted to individual tiff volumes using Fiji.

### Automated lineage tracing and manual curation

Individual TIFF volumes were cropped to 512×512 pixel square images, converted to 32-bit floating point values and processed slice-by-slice with embGAN. embGAN output slices were reassembled into multipage TIFF stacks using Fiji and processed using StarryNite and the resulting tracking data were curated using AceTree. Cell identities and division times were checked by hand and corrected if necessary. Parameter files for StarryNite processing are included in the raw data archive on DataDryad. Cells were individually tracked confirming the timing of cell divisions up until the generational targets for each sub-lineage. Each lineage was manually curated to the 100-cell stage (the sixth generation of the AB lineage, the fourth of the MS lineage, the third of the E and C lineages, and the second of the D lineage). Several cells were omitted from analysis across all embryos since they were unable to be tracked in certain embryos, which were MSpppp, MSpppa, ABalaapa, ABarppaap, ABprpaap, ABprppaa, and MSppaa. The lineage was truncated prior to P4 division, excluding P4, Z2, and Z3 from analysis. Two embryos from the CB4856 strain were excluded from analysis, because they had a spontaneous developmental pause in their E and D lineages. Although previous work has indicated that developmental pauses in the embryo could be due to transient factors such as hypoxia^34^, it is unclear what might have caused this pause in just two lineages. Subsequent attempts to reproduce this phenomenon were unsuccessful.

### Branch distance

The branch distance was developed as a metric enabling calculations of distance between cell lineages, where a quantitative measure has been mapped onto individual cells^28^. This metric was demonstrated to contain particular utility for the comparison of *C. elegans* cell-cycle durations during early embryonic development, as its invariant lineage enables precise cell matching between individuals. Here, we calculated the branch distance in a pairwise manner between lineage trees where the corresponding cell-cycle time was matched to each cell. The Euclidean norm was calculated on the differences between matching cell-cycle times for all cells in two embryos, resulting in a single value measuring the relative dissimilarity in cell-cycle timing between two lineages. Two versions of this metric were developed, the union and intersection branch distance, where the union branch distance accounts for differences in tree topology, and the intersection branch distance does not. In this case, as we compared development of different wild strains of *C. elegans*, all trees were topologically identical due to their invariant lineage under normal conditions. Throughout, we refer to this distance as just the branch distance, because the union and intersection branch distance were functionally identical in this case.

### Permutation test framework

To formalize observed inter-strain differences in cell-cycle timing distributions, we developed a non-parametric permutation-based statistical framework. In this test, all strains were iterated through in a pair-wise manner. For each strain-to-strain comparison the mean branch distance within each strain was calculated, averaged between the two values, and then subtracted from the mean inter-strain branch distance. The embryos were then shuffled between strains, these values were recalculated, and a greater inter - intra branch distance was counted as a positive result. P values were calculated by dividing the number of positive results over the total number of permutation iterations for each pairwise comparison, and then underwent Benjamini-Hochberg correction after pooling across all strain and lineage comparisons for a given version of the branch distance (normal, parallel, perpendicular). Q values thresholded at or below 0.05 were then aggregated to understand which lineages and/or strains displayed the greatest number of differences. Tabulation of these differences was performed by summing up within each strain and lineage, with a maximum of seven if a given lineage within a strain was different from all other seven strains. These results were then summed across all lineages within a strain to measure strain divergence and across all strains within a given lineage to measure lineage divergence.

### Branch distance decomposition

Although the branch distance has served as our aggregate measure of lineage-aligned timing differences between embryos, the metric conflates two qualitatively distinct sources of variance. Strains exhibit apparent clock-rate differences that are somewhat consistent across the entire lineage, meaning that some portion of the distances calculated previously may reflect global scaling in cell-cycle times between embryos rather than localized rearrangements of developmental timing within lineages. Critically, different mechanisms might drive these two distinct sources of variation. Correlated changes in timing across lineages could be driven by biophysical mechanisms, metabolism, or maternal load for instance, and residual variation could arise due to local causes such as signaling, fate-specific behaviors, or due to noise alone. We therefore sought to decompose the branch distance into a global scaling component and a residual capturing all other lineage-aligned variation.

To accomplish this decomposition, we represented the cell cycles for a given embryo as a vector, where every dimension corresponds to the cell-cycle duration of a particular cell (Figure 5A). For any pair of embryos, the branch distance is the Euclidean norm of the difference between the two cell-cycle vectors, computed over the shared set of measured cells.

**Figure 5:**
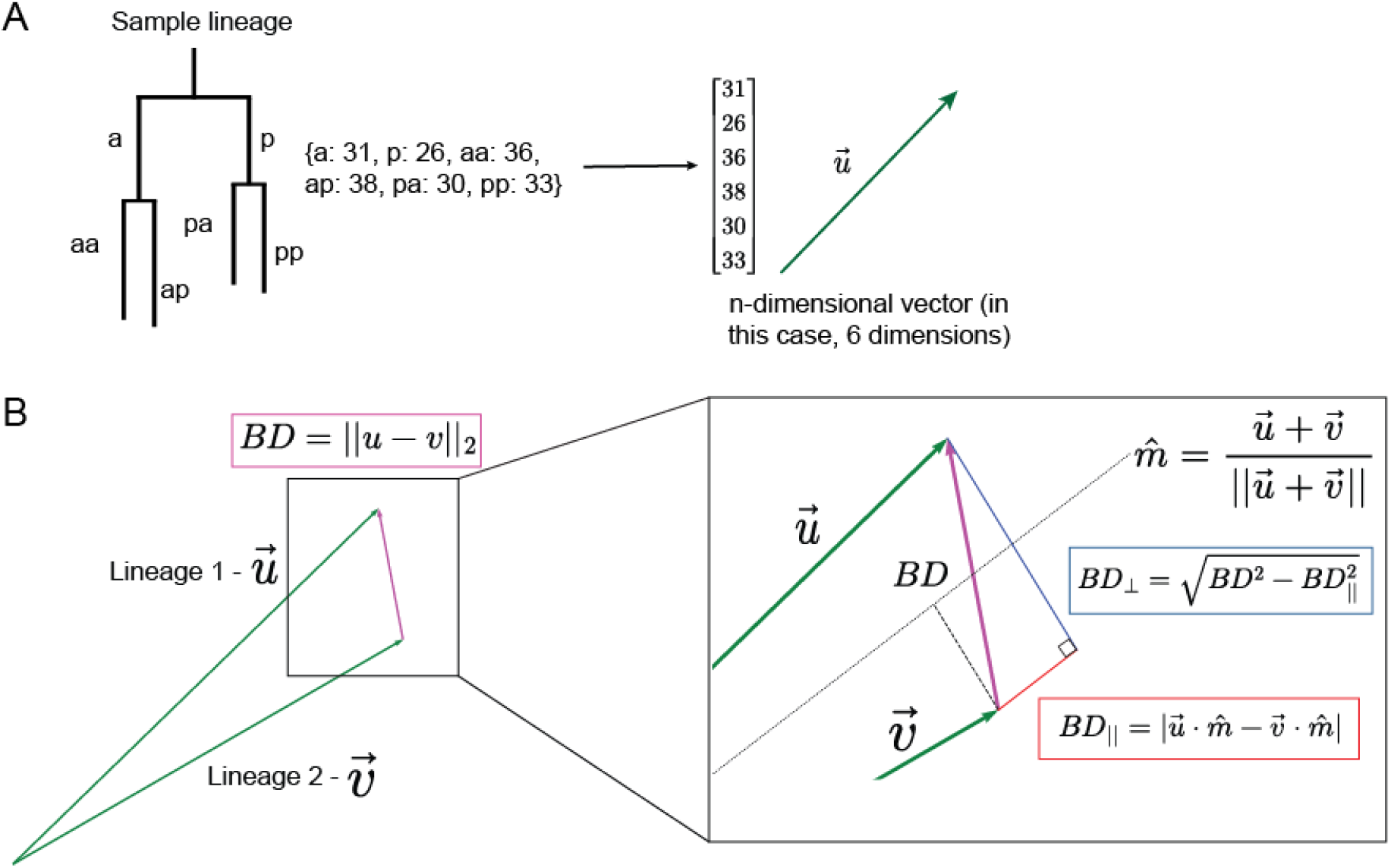
A: Diagrammatic representation of the conversion of a lineage tree to its corresponding n-dimensional vector. B: Graphical representation of the decomposition of the branch distance into its parallel and perpendicular components. This is calculated by first calculating the mean unit vector between any pair of lineage vectors. The absolute difference between the dot product of each lineage vector and the mean vector becomes the parallel component, and the perpendicular component is calculated by taking the root-squared difference between the branch distance and the calculated parallel component of the branch distance.

We then decomposed this difference vector into two components: a parallel component aligned with the bisecting vector of the two embryos’ cell-cycle vectors, that is the vector 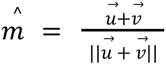 that lies symmetrically between them, and a perpendicular residual component. Geometrically, the parallel direction captures uniform scaling of one embryo’s developmental tempo relative to the other: if embryo A’s cell cycles are uniformly α-fold longer than embryo B’s, the difference vector lies exactly along the bisector and the perpendicular residual is zero. The bisector was chosen rather than the dataset mean or a single reference because it is symmetric in 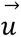 and 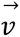 - guaranteeing that 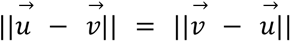 under decomposition, and because it recovers pure global scaling without privileging any particular reference. Deviations from pure uniform scaling, whether a single cell is cycling at an anomalous rate, or a coordinated shift confined to a sub-branch of a lineage, would project onto the perpendicular component itself.

The parallel component thus captures global linear scaling in developmental timing between embryos, whereas the residual component represents lineage-specific deviations and any other non-scaling differences in developmental structure. By construction, the squared branch distance partitions exactly: 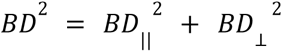. Applied to whole embryos, the decomposition separates pairwise differences in global developmental rate from pairwise differences in lineage structure. Applied lineage by lineage, it asks whether the strain-specific timing differences arise from uniform scaling within each lineage or from non-scaling or cell or branch-specific rearrangements. We tested both components by the permutation framework introduced above and quantified the frequency of significant pairwise inter-strain differences for each.

## Data Availability Statement

Cell-cycle timing measurements for all embryos in this study and the analysis code that generates every figure are available for access (DOI: 10.5281/zenodo.23813615).

Wild strain images are deposited as 8-bit TIFFs and are accompanied by the raw and curated AceTree-format lineage output in the Dryad Digital Repository (DOI pending). The embGAN-generated outputs were omitted for storage efficiency; they can be regenerated from the 8-bit images using embGAN^27^(DOI: 10.5281/zenodo.10535870).

GitHub Copilot was used to assist the writing and editing of Python code for this manuscript. All suggested code was checked for accuracy, and the authors take full responsibility for this content.

## Acknowledgements

This work was supported by the National Institutes of Health (R35GM151199 to PKS.; NIH R01GM143378 to EJD) and the National Science Foundation (NSF 2524223 to ECA.).

## Supplementary Material

**Figure S1.**
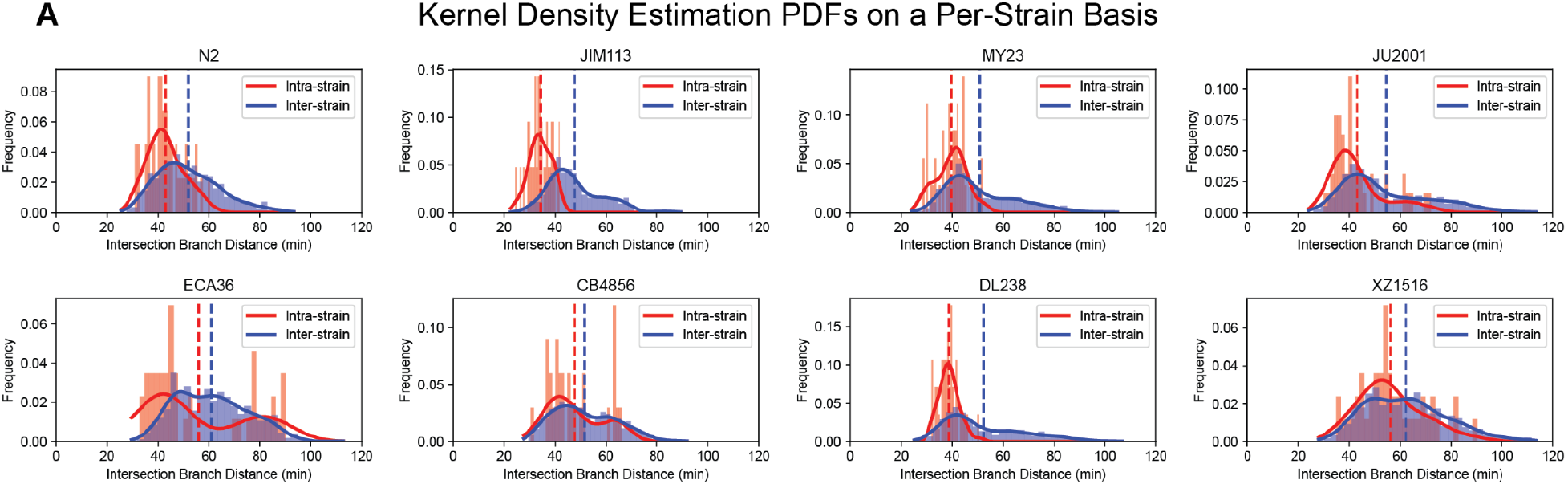
Distribution of within-strain (red) and between strain (blue) branch distances across all eight strains. Dashed lines show distribution means, histograms show observed distributions, and solid lines show Gaussian kernel density estimator fits.

**Table S1.**
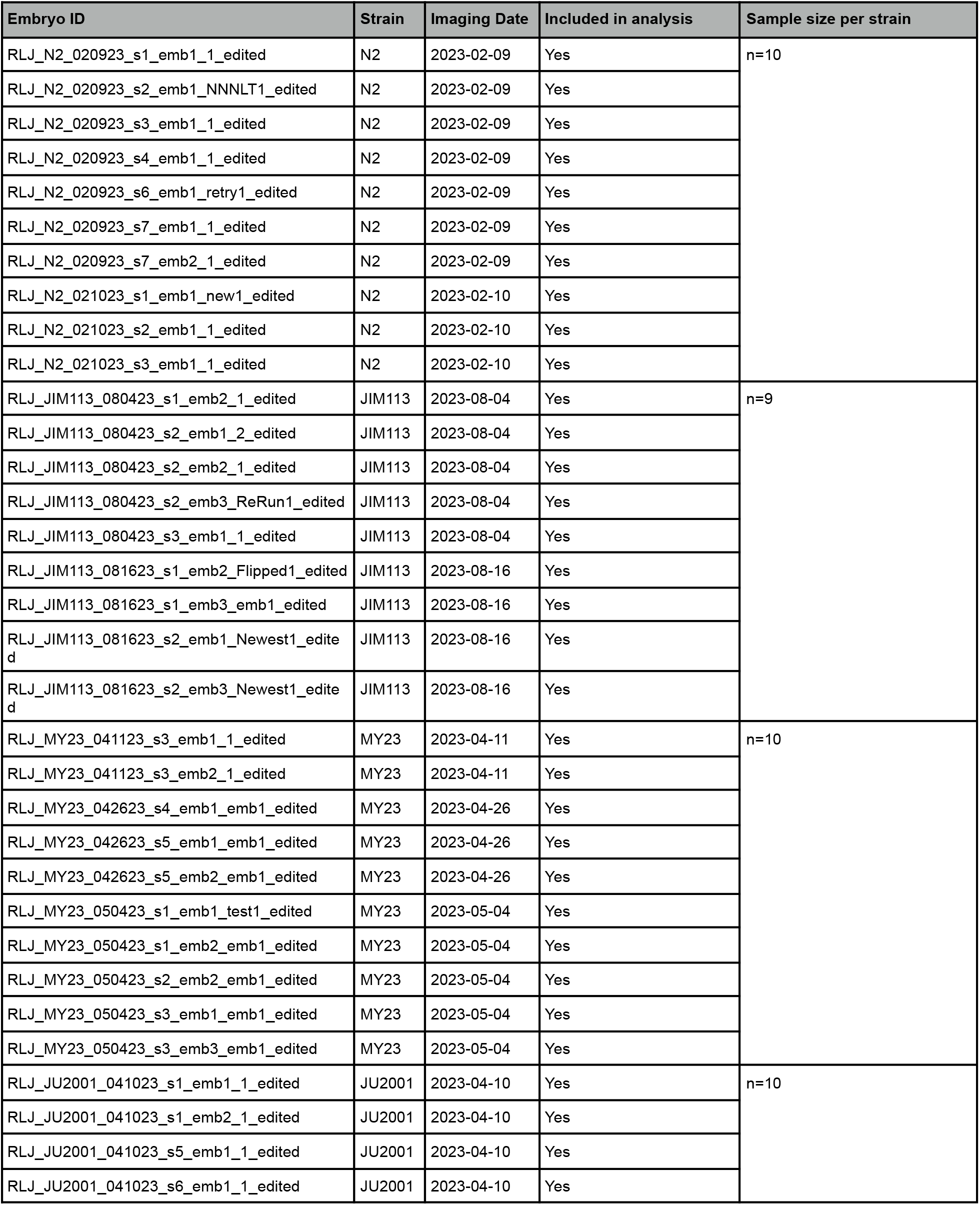
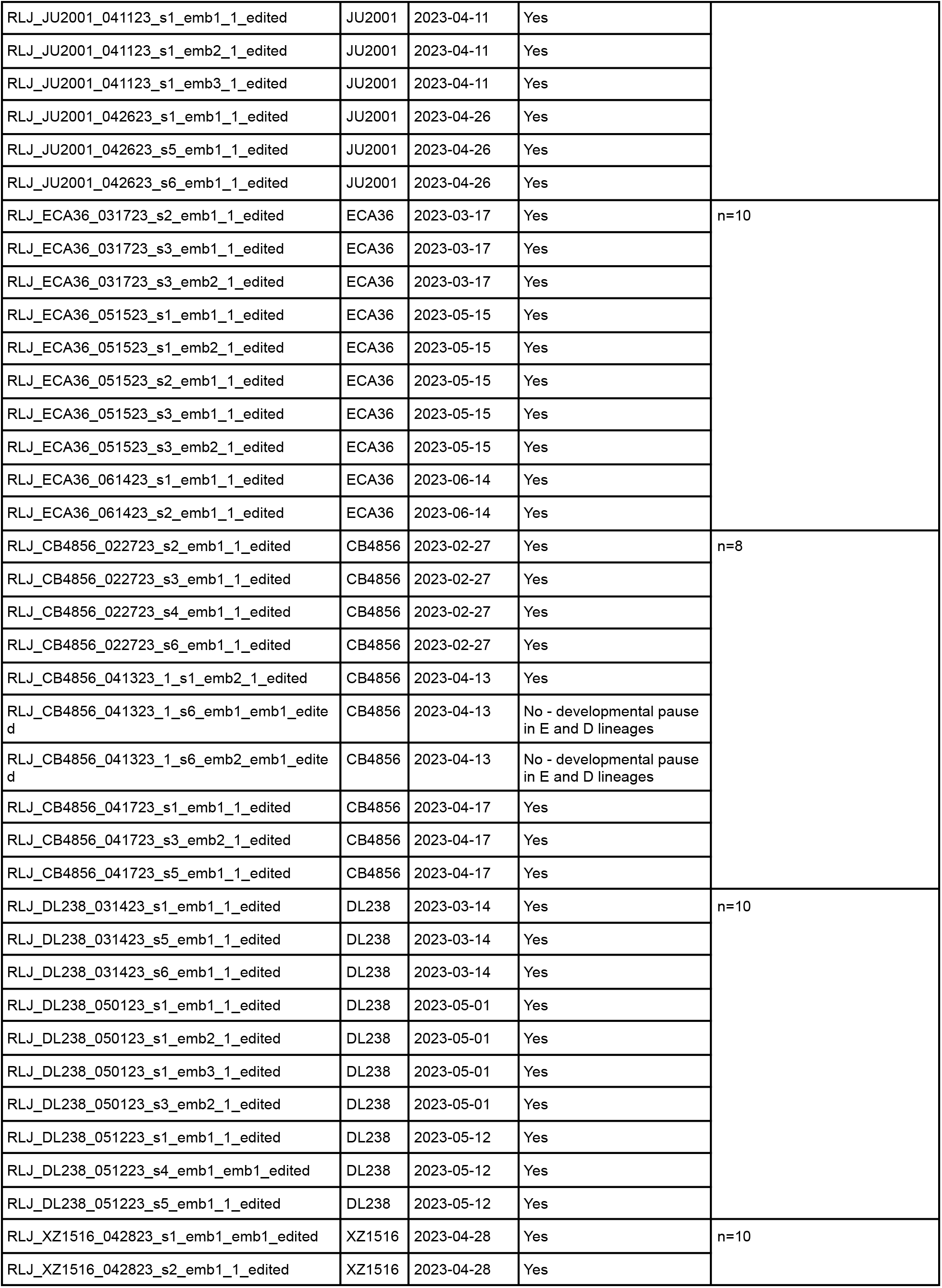
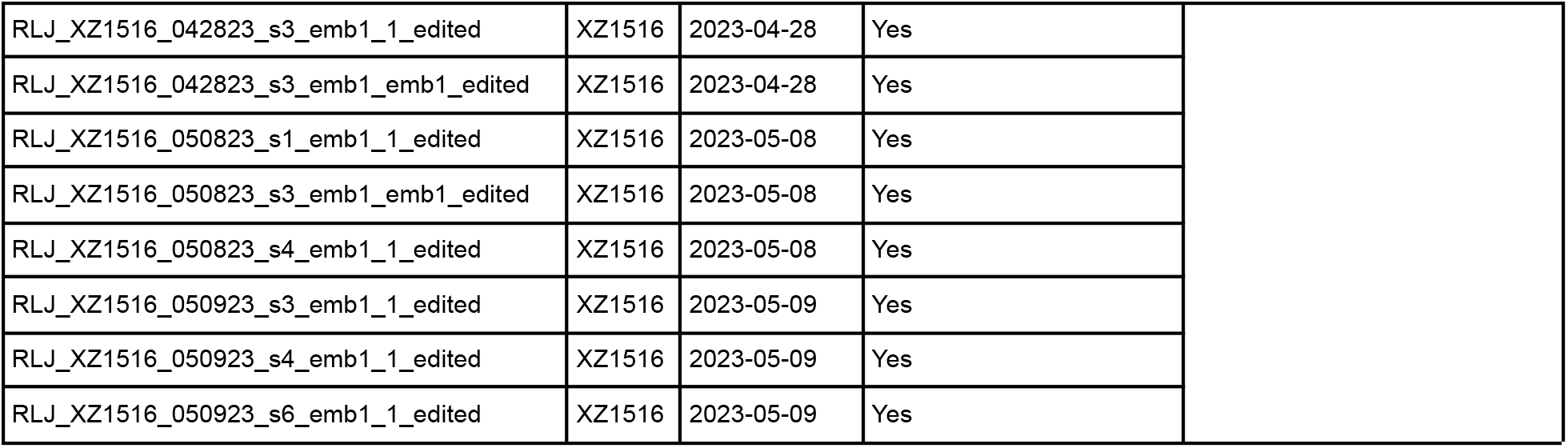
Embryos in dataset labelled with their ID, strain label, imaging date, inclusion in analysis, and sample size.

| Embryo ID | Strain | Imaging Date | Included in analysis | Sample size per strain |
|---|---|---|---|---|
| RLJ_N2_020923_s1_emb1_1_edited | N2 | 2023-02-09 | Yes | n=10 |
| RLJ_N2_020923_s2_emb1_NNNLT1_edited | N2 | 2023-02-09 | Yes |  |
| RLJ_N2_020923_s3_emb1_1_edited | N2 | 2023-02-09 | Yes |  |
| RLJ_N2_020923_s4_emb1_1_edited | N2 | 2023-02-09 | Yes |  |
| RLJ_N2_020923_s6_emb1_retry1_edited | N2 | 2023-02-09 | Yes |  |
| RLJ_N2_020923_s7_emb1_1_edited | N2 | 2023-02-09 | Yes |  |
| RLJ_N2_020923_s7_emb2_1_edited | N2 | 2023-02-09 | Yes |  |
| RLJ_N2_021023_s1_emb1_new1_edited | N2 | 2023-02-10 | Yes |  |
| RLJ_N2_021023_s2_emb1_1_edited | N2 | 2023-02-10 | Yes |  |
| RLJ_N2_021023_s3_emb1_1_edited | N2 | 2023-02-10 | Yes |  |
| RLJ_JIM113_080423_s1_emb2_1_edited | JIM113 | 2023-08-04 | Yes | n=9 |
| RLJ_JIM113_080423_s2_emb1_2_edited | JIM113 | 2023-08-04 | Yes |  |
| RLJ_JIM113_080423_s2_emb2_1_edited | JIM113 | 2023-08-04 | Yes |  |
| RLJ_JIM113_080423_s2_emb3_ReRun1_edited | JIM113 | 2023-08-04 | Yes |  |
| RLJ_JIM113_080423_s3_emb1_1_edited | JIM113 | 2023-08-04 | Yes |  |
| RLJ_JIM113_081623_s1_emb2_Flipped1_edited | JIM113 | 2023-08-16 | Yes |  |
| RLJ_JIM113_081623_s1_emb3_emb1_edited | JIM113 | 2023-08-16 | Yes |  |
| RLJ_JIM113_081623_s2_emb1_Newest1_edited | JIM113 | 2023-08-16 | Yes |  |
| RLJ_JIM113_081623_s2_emb3_Newest1_edited | JIM113 | 2023-08-16 | Yes |  |
| RLJ_MY23_041123_s3_emb1_1_edited | MY23 | 2023-04-11 | Yes | n=10 |
| RLJ_MY23_041123_s3_emb2_1_edited | MY23 | 2023-04-11 | Yes |  |
| RLJ_MY23_042623_s4_emb1_emb1_edited | MY23 | 2023-04-26 | Yes |  |
| RLJ_MY23_042623_s5_emb1_emb1_edited | MY23 | 2023-04-26 | Yes |  |
| RLJ_MY23_042623_s5_emb2_emb1_edited | MY23 | 2023-04-26 | Yes |  |
| RLJ_MY23_050423_s1_emb1_test1_edited | MY23 | 2023-05-04 | Yes |  |
| RLJ_MY23_050423_s1_emb2_emb1_edited | MY23 | 2023-05-04 | Yes |  |
| RLJ_MY23_050423_s2_emb2_emb1_edited | MY23 | 2023-05-04 | Yes |  |
| RLJ_MY23_050423_s3_emb1_emb1_edited | MY23 | 2023-05-04 | Yes |  |
| RLJ_MY23_050423_s3_emb3_emb1_edited | MY23 | 2023-05-04 | Yes |  |
| RLJ_JU2001_041023_s1_emb1_1_edited | JU2001 | 2023-04-10 | Yes | n=10 |
| RLJ_JU2001_041023_s1_emb2_1_edited | JU2001 | 2023-04-10 | Yes |  |
| RLJ_JU2001_041023_s5_emb1_1_edited | JU2001 | 2023-04-10 | Yes |  |
| RLJ_JU2001_041023_s6_emb1_1_edited | JU2001 | 2023-04-10 | Yes |  |
| RLJ_JU2001_041123_s1_emb1_1_edited | JU2001 | 2023-04-11 | Yes |  |
| RLJ_JU2001_041123_s1_emb2_1_edited | JU2001 | 2023-04-11 | Yes |  |
| RLJ_JU2001_041123_s1_emb3_1_edited | JU2001 | 2023-04-11 | Yes |  |
| RLJ_JU2001_042623_s1_emb1_1_edited | JU2001 | 2023-04-26 | Yes |  |
| RLJ_JU2001_042623_s5_emb1_1_edited | JU2001 | 2023-04-26 | Yes |  |
| RLJ_JU2001_042623_s6_emb1_1_edited | JU2001 | 2023-04-26 | Yes |  |
| RLJ_ECA36_031723_s2_emb1_1_edited | ECA36 | 2023-03-17 | Yes | n=10 |
| RLJ_ECA36_031723_s3_emb1_1_edited | ECA36 | 2023-03-17 | Yes |  |
| RLJ_ECA36_031723_s3_emb2_1_edited | ECA36 | 2023-03-17 | Yes |  |
| RLJ_ECA36_051523_s1_emb1_1_edited | ECA36 | 2023-05-15 | Yes |  |
| RLJ_ECA36_051523_s1_emb2_1_edited | ECA36 | 2023-05-15 | Yes |  |
| RLJ_ECA36_051523_s2_emb1_1_edited | ECA36 | 2023-05-15 | Yes |  |
| RLJ_ECA36_051523_s3_emb1_1_edited | ECA36 | 2023-05-15 | Yes |  |
| RLJ_ECA36_051523_s3_emb2_1_edited | ECA36 | 2023-05-15 | Yes |  |
| RLJ_ECA36_061423_s1_emb1_1_edited | ECA36 | 2023-06-14 | Yes |  |
| RLJ_ECA36_061423_s2_emb1_1_edited | ECA36 | 2023-06-14 | Yes |  |
| RLJ_CB4856_022723_s2_emb1_1_edited | CB4856 | 2023-02-27 | Yes | n=8 |
| RLJ_CB4856_022723_s3_emb1_1_edited | CB4856 | 2023-02-27 | Yes |  |
| RLJ_CB4856_022723_s4_emb1_1_edited | CB4856 | 2023-02-27 | Yes |  |
| RLJ_CB4856_022723_s6_emb1_1_edited | CB4856 | 2023-02-27 | Yes |  |
| RLJ_CB4856_041323_1_s1_emb2_1_edited | CB4856 | 2023-04-13 | Yes |  |
| RLJ_CB4856_041323_1_s6_emb1_emb1_edited | CB4856 | 2023-04-13 | No - developmental pause in E and D lineages |  |
| RLJ_CB4856_041323_1_s6_emb2_emb1_edited | CB4856 | 2023-04-13 | No - developmental pause in E and D lineages |  |
| RLJ_CB4856_041723_s1_emb1_1_edited | CB4856 | 2023-04-17 | Yes |  |
| RLJ_CB4856_041723_s3_emb2_1_edited | CB4856 | 2023-04-17 | Yes |  |
| RLJ_CB4856_041723_s5_emb1_1_edited | CB4856 | 2023-04-17 | Yes |  |
| RLJ_DL238_031423_s1_emb1_1_edited | DL238 | 2023-03-14 | Yes | n=10 |
| RLJ_DL238_031423_s5_emb1_1_edited | DL238 | 2023-03-14 | Yes |  |
| RLJ_DL238_031423_s6_emb1_1_edited | DL238 | 2023-03-14 | Yes |  |
| RLJ_DL238_050123_s1_emb1_1_edited | DL238 | 2023-05-01 | Yes |  |
| RLJ_DL238_050123_s1_emb2_1_edited | DL238 | 2023-05-01 | Yes |  |
| RLJ_DL238_050123_s1_emb3_1_edited | DL238 | 2023-05-01 | Yes |  |
| RLJ_DL238_050123_s3_emb2_1_edited | DL238 | 2023-05-01 | Yes |  |
| RLJ_DL238_051223_s1_emb1_1_edited | DL238 | 2023-05-12 | Yes |  |
| RLJ_DL238_051223_s4_emb1_emb1_edited | DL238 | 2023-05-12 | Yes |  |
| RLJ_DL238_051223_s5_emb1_1_edited | DL238 | 2023-05-12 | Yes |  |
| RLJ_XZ1516_042823_s1_emb1_emb1_edited | XZ1516 | 2023-04-28 | Yes | n=10 |
| RLJ_XZ1516_042823_s2_emb1_1_edited | XZ1516 | 2023-04-28 | Yes |  |
| RLJ_XZ1516_042823_s3_emb1_1_edited | XZ1516 | 2023-04-28 | Yes |  |
| RLJ_XZ1516_042823_s3_emb1_emb1_edited | XZ1516 | 2023-04-28 | Yes |  |
| RLJ_XZ1516_050823_s1_emb1_1_edited | XZ1516 | 2023-05-08 | Yes |  |
| RLJ_XZ1516_050823_s3_emb1_emb1_edited | XZ1516 | 2023-05-08 | Yes |  |
| RLJ_XZ1516_050823_s4_emb1_1_edited | XZ1516 | 2023-05-08 | Yes |  |
| RLJ_XZ1516_050923_s3_emb1_1_edited | XZ1516 | 2023-05-09 | Yes |  |
| RLJ_XZ1516_050923_s4_emb1_1_edited | XZ1516 | 2023-05-09 | Yes |  |
| RLJ_XZ1516_050923_s6_emb1_1_edited | XZ1516 | 2023-05-09 | Yes |  |

